# Hippocampo-cortical circuits for selective memory encoding, routing, and replay

**DOI:** 10.1101/2022.09.25.509420

**Authors:** Ryan E. Harvey, Heath L. Robinson, Can Liu, Azahara Oliva, Antonio Fernandez-Ruiz

**Affiliations:** Department of Neurobiology & Behavior, Cornell University, Ithaca, USA

## Abstract

Traditionally considered a homogeneous cell type, hippocampal pyramidal cells have been recently shown to be highly diverse. However, how this cellular diversity relates to the different hippocampal network computations that support memory-guided behavior is not yet known. We discovered that the anatomical identity of pyramidal cells is a major organizing principle of CA1 assembly dynamics, the emergence of memory replay, and cortical projection patterns. Segregated pyramidal cell subpopulations encoded trajectory and choice-specific information or tracked changes in reward configuration respectively, and selectively routed these representations to different cortical targets. Furthermore, distinct hippocampo-cortical assemblies coordinated the reactivation of complementary memory representations. These findings reveal the existence of specialized hippocampo-cortical subcircuits and provide a cellular mechanism that supports the computational flexibility and memory capacities of these structures.

---

Neural ensemble interactions in the hippocampus and associated cortical structures support learning and memory. Ensemble reactivation during sharp-wave ripples (SWRs), generated in the CA1 area of the hippocampus, broadcast memory representations to the rest of the brain, a process that supports consolidation, retrieval, planning, prospective, and retrospective coding [1–8]. Despite this multiplicity of functions, the hippocampus has been traditionally conceived as being composed of functionally homogenous principal cells in each of its subregions. Contrary to this notion, recent studies have highlighted that hippocampal pyramidal cells are indeed very diverse in their molecular, developmental, anatomical, and physiological characteristics [9–17]. To date, it is not yet understood how this cellular diversity relates to the different computational operations and functional correlates displayed by the hippocampus during navigation and learning. To investigate how hippocampal cellular heterogeneity supports diverse memory functions, we examined the dynamic organization and behavioral correlates of large neuronal ensembles in the hippocampus and two of its major targets, the medial entorhinal (MEC) and prefrontal (PFC) cortices in rats during memory tasks and sleep.

CA1 principal cell heterogeneity strongly correlates with their radial (depth) position within the pyramidal layer [9, 14–16, 19–26]. To map this neuronal diversity, we examined a large number of simultaneously recorded cells using high-density silicon probes that span the radial dorsal CA1 axis in rats (Fig. 1A-B and Fig. S1; n = 9,838 cells in 49 rats). The stereotypical depth profile of SWRs in CA1 allowed us to divide the pyramidal layer into deep, middle, and superficial sublayers and classify all recorded cells accordingly (Fig. 1B and Fig. S1 [16]). In agreement with previous reports, we found that deep (CA1deep) and superficial (CA1sup) pyramidal cells differ in their physiological properties (Fig. S1; [16, 17, 22, 23, 26]).

**Figure 1:**
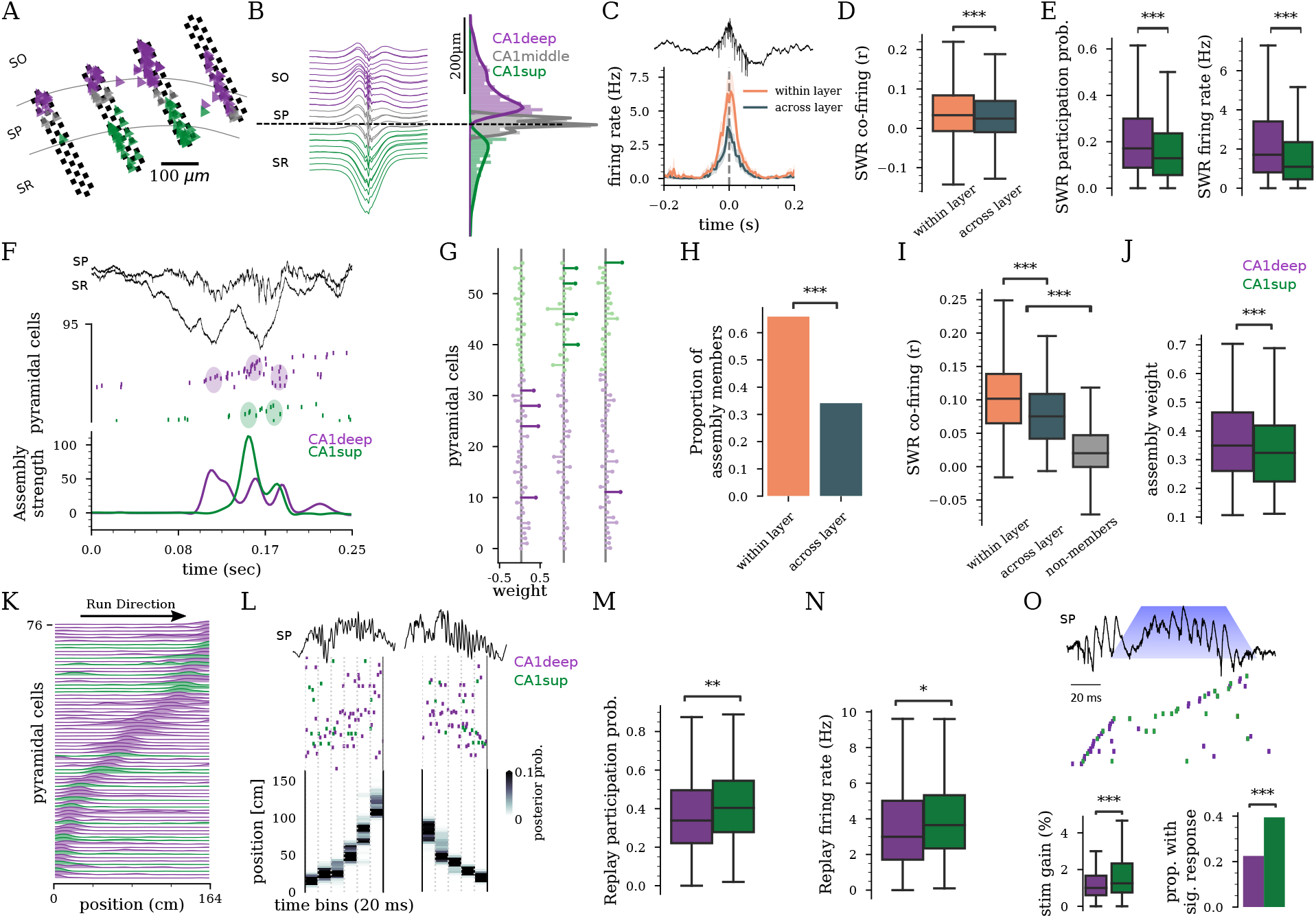
Anatomical organization of hippocampal assembly dynamics and memory replay. **A)** Schematic of 128-channel probe in CA1 sublayers: stratum oriens (SO), pyramidal layer (SP), stratum radiatum (SR) with overlay of pyramidal cell’s (triangles) from deep (purple) to superficial (green). **B)** Left: Average SWR profile. Right: distribution of CA1deep (3,658), CA1middle (2,706), and CA1sup (1,946) pyramidal cells in 49 rats. **C)** Example SWR cross-correlogram of cells within and across sublayers. **D)** Within and across sublayer SWR co-firing (*P* < 2 × 10^-16^, Linear mixed-effects model (LME)). **E)** SWR participation probability and firing rate for CA1deep and CA1sup (*P* = 2.33 × 10^-6^ / *P* = 1.51 × 10^-8^, LME). **F)** SWR (top) with cell firing (middle) and activation of assemblies (bottom). **G)** Example assembly weights. Length indicates weight. Assembly members are darker. **H)** Assembly member composition from within and across layers (n=387 assemblies, *P* = 2.56 × 10^-19^, χ^2^). **I)** Within and across layer SWR co-firing of assembly members and non-members (*P* = 2.48 × 10^-3^ / *P* < 2 × 10^-16^, LME). **J)** Contribution to assembly (weight) for CA1deep and CA1sup (*P* = 7.23 × 10^-5^, LME). **K)** Place fields on novel linear track. **L)** Example raster (middle) during a SWR (top) and decoded animal position (bottom) for replay events. **M)** Probability of participation in replay (*P* = 2.14 × 10^-13^, LME, n=671 cells, 12 rats). **N)** Firing rate in replay (*P* = .044, LME) **O)** Top: optogenetically prolonged SWR and cell sequence in memory task [18]. Bottom left: Firing rate increase in prolonged SWR vs. baseline (n=467 CA1deep / 289 CA1sup, 5 rats, *P* = 2.52 × 10^-4^, LME). Bottom right: Proportion of CA1deep and CA1sup recruited by stimulation (n=1074 events, *P* = 5.84 × 10^-7^, χ^2)^.

Contrary to many cortical areas, the hippocampus has been traditionally conceived as having a salt-and-pepper organization, where the correlated firing dynamics of its principal cells are independent of their anatomical location [27–29]. To investigate whether CA1deep and CA1sup cells are functionally correlated, we first analyzed their firing during SWRs, as these are events of enhanced network synchrony [30]. We found that pyramidal cells from the same sublayer were more likely to fire together during SWRs compared to cells from different sublayers (Fig. 1D). Although both CA1deep and CA1sup cells were strongly recruited during SWRs, CA1deep had a higher participation probability and fired more during SWRs compared to CA1sup (Fig. 1E). The synchronous co-activation of pyramidal cells that form functional “assemblies” has been proposed as the building block of hippocampal and cortical representations [28, 31–33]. We analyzed CA1 pyramidal cell assemblies using established methods, and found a strong anatomical bias in their cellular composition (Fig. 1F-H). In two-thirds of assemblies, all members belonged to the same sublayer (Fig. 1G-H). Pairs of assembly members from the same sublayer co-fired during SWRs more often than members of different sublayers or non-member cells (Fig. 1I). Overall, CA1deep contributed more to assembly activation than CA1sup (Fig. 1J). We found similar results when applying these analyses to theta activity during behavior, rather than SWRs (Fig. S2), suggesting that a higher synchronization between cells from the same sublayer is a general principle of hippocampal network dynamics.

To directly assess the contribution of CA1deep and CA1sup to memory, we analyzed SWR-associated replay during the exploration of novel mazes and subsequent sleep sessions. We trained a Bayesian decoding algorithm on place cell activity on the linear track (Fig. 1K and Fig. S3) and applied it to all SWR-associated spiking sequences to identify replay events (Fig. 1L and Fig. S3). Given the higher SWR participation of CA1deep (Fig. 1E,J), we expected CA1deep cells to also dominate replay events. Surprisingly, it was CA1sup cells that were specifically recruited and fired more during replay events than CA1deep (Fig. 1M-N). This preferential recruitment of CA1sup cells led to the hypothesis that CA1sup cells are preferentially involved in memory processes. Given that we have previously shown that optogenetic prolongation of SWRs during learning improves rats’ performance in a delayed alternation memory task [18], we analyzed whether this improvement was mediated by CA1sup cells. Indeed, our optogenetic manipulation prolonged ongoing SWRs by preferentially recruiting CA1sup cells, rather than CA1deep cells, into the ongoing spiking sequence (Fig. 1O).

CA1 broadcasts hippocampal memory representations to multiple areas of the brain during SWRs [4, 6, 7, 34–38]. To investigate whether CA1deep and CA1sup selectively route information to different downstream target areas, we focused on two hippocampal main output regions implicated in memory-guided behavior: MEC and PFC. First, we examined dorsal hippocampal projections to these regions using fluorescent retrograde tracers (Fig. 2A and Fig. S4). Most monosynaptic inputs from CA1 to PFC originated in the deep sublayer, while most inputs to MEC originated from superficial pyramidal cells (Fig. 2B-C and Fig S4). We found a larger proportion of MEC than PFC projecting pyramidal cells and only a small minority of dual-projecting ones (Fig. 2C), suggesting these subpopulations form largely parallel circuits.

**Figure 2:**
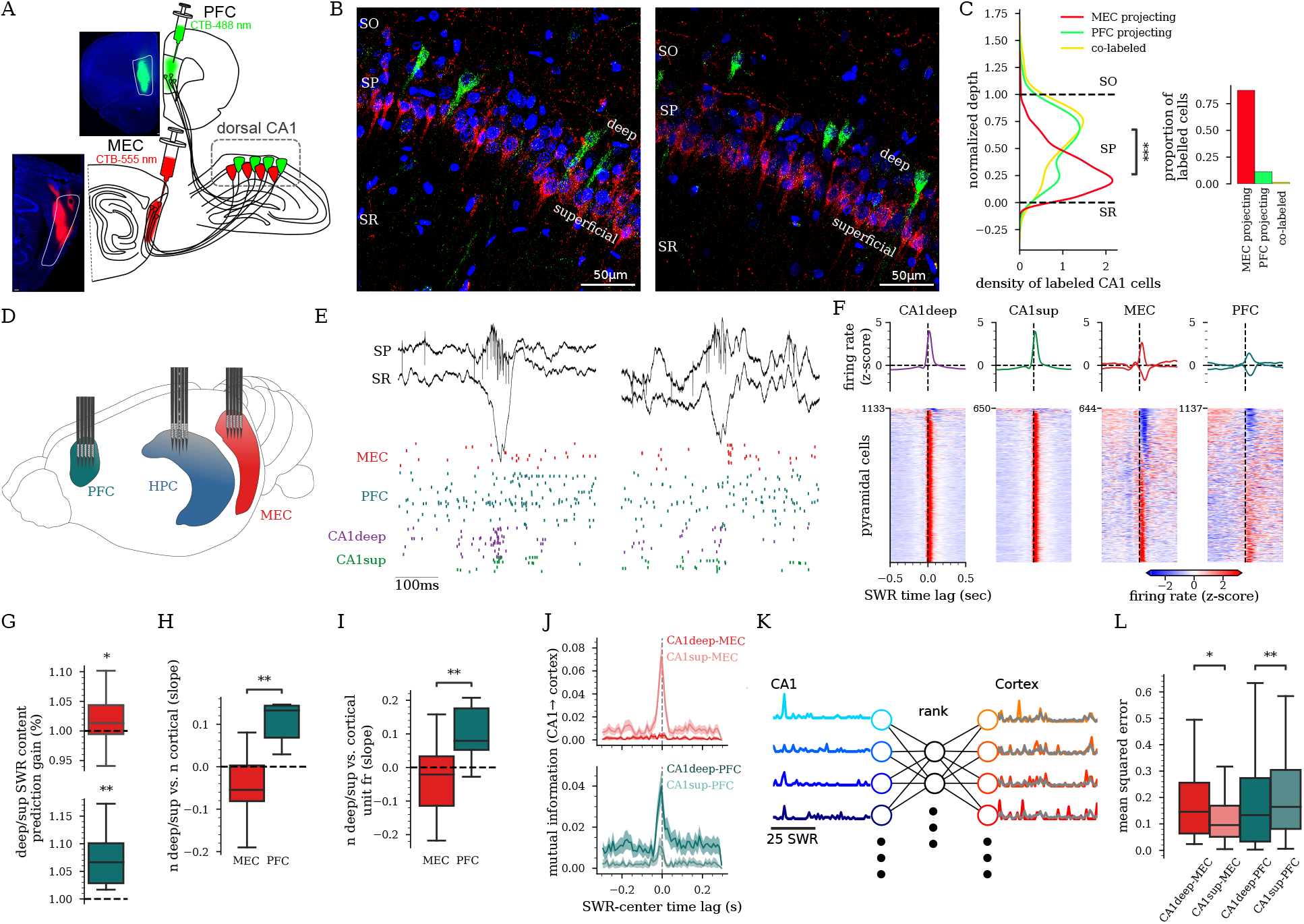
CA1deep and CA1sup route information to different downstream cortical areas. **A)** Green and red fluorescent retrograde tracers (Alexa Fluor-CTB conjugates) were injected in PFC and MEC to label hippocampal projecting cells. **B)** Representative histology from the same animal as A with CA1deep-PFC projecting (green) and CA1sup-MEC projecting (red) cells in dorsal hippocampus. Blue: DAPI, Red: CTB-555, Green: CTB488. **C)** Quantification of soma location for labeled cells (n=279 PFC projecting / 2,167 MEC projecting / 31 co-labeled cells in 3 rats, *P* < 2 × 10^-16^, rank-sum test, PFC-projecting vs. MEC-projecting depth). **D)** Recording schematic. **E)** Example unit responses (bottom) to SWR (top) in each region. **F)** Peri-SWR firing rate for each region. Average responses (±95% CI) (top) (positive and negative responses separated for cortex) and individual units (below). **G)** Prediction gain of the ratio of (CA1deep/CA1sup) for cells active in SWRs calculated from cortical responses vs. shuffle distributions (MEC: *P* = 0.043, PFC: *P* = 2.3 × 10^-3^, t-test). **H-I)** Regression slopes between SWR relative content of CA1deep vs. CA1sup cells **H)** and number of cortical units that responded (*P* = 1.534 × 10^-3^, rank-sum), and **I)** cortical firing rate responses in SWRs (*P* = 4.35 × 10^-3^, rank-sum). **J)** Examples of peri-SWR mutual information between cross-region cell pairs (mean±95%CI). **K)** Schematic of reduced rank regression used to predict neural activity in cortex from neural activity in CA1 during SWRs. **L)** Mean squared error of predictions of cortical SWR responses from CA1 activity patterns using reduced rank regression (*P* = 0.04 for CA1deep/sup versus MEC, *P* = 1.33 × 10^-4^ for CA1deep/sup vs. PFC, LME).

To analyze the functional implication of our anatomical findings, we performed simultaneous ensemble recordings in CA1, PFC, and MEC (Fig. 2D and Fig. S5). Neurons in the three regions were strongly entrained by SWRs (Fig. 2E-F; [34, 39]). Using a linear model, we found that cortical ensembles discriminate the anatomical distribution of CA1 cells participating in SWRs (Fig. 2G). Next, we examined the magnitude of cortical activity during SWRs as a function of their relative ‘cell content’. A larger proportion of MEC cells were recruited and fired more in response to SWRs with predominantly CA1sup content (Fig. 2H-I). Likewise, a larger proportion of PFC cells were recruited and fired more in response to SWRs with predominant CA1deep content (Fig. 2H-I). We then examined if MEC and PFC ensembles preferentially responded to the firing patterns of deep and superficial cells. We found significantly higher mutual information between the activity of CA1sup and MEC cells and between CA1deep and PFC cells during SWRs (Fig. 2J; CA1sup and MEC *P* = 0.039, CA1deep and PFC *P* = 0.045, LME). To directly test the extent to which hippocampal ensemble activity predicted cortical population responses during individual SWRs, we performed reduced rank regression (Fig. 2K). We obtained better predictions of MEC responses from CA1sup SWR patterns and of PFC responses from CA1deep patterns (Fig. 2L), in agreement with our anatomical tracing results.

To determine if CA1deep and CA1sup encode different task-related information and selectivity route to PFC and MEC, we analyzed their activity during behavior. First, we first examined place cell dynamics across multiple environments on the same day (Fig. 3A and Fig. S6). Similar to previous reports, we found a larger proportion of place cells among CA1deep compared to CA1sup (68% vs. 50% *P* < .0001 χ^2^, n = 1,645 place cells in 19 rats, Fig. S6; [16, 17, 23, 24]). CA1deep cells were more likely to develop fields in multiple environments than CA1sup cells, while the firing of CA1sup cells was more informative about specific environment identity (Fig. 3A and Fig. S6).

**Figure 3:**
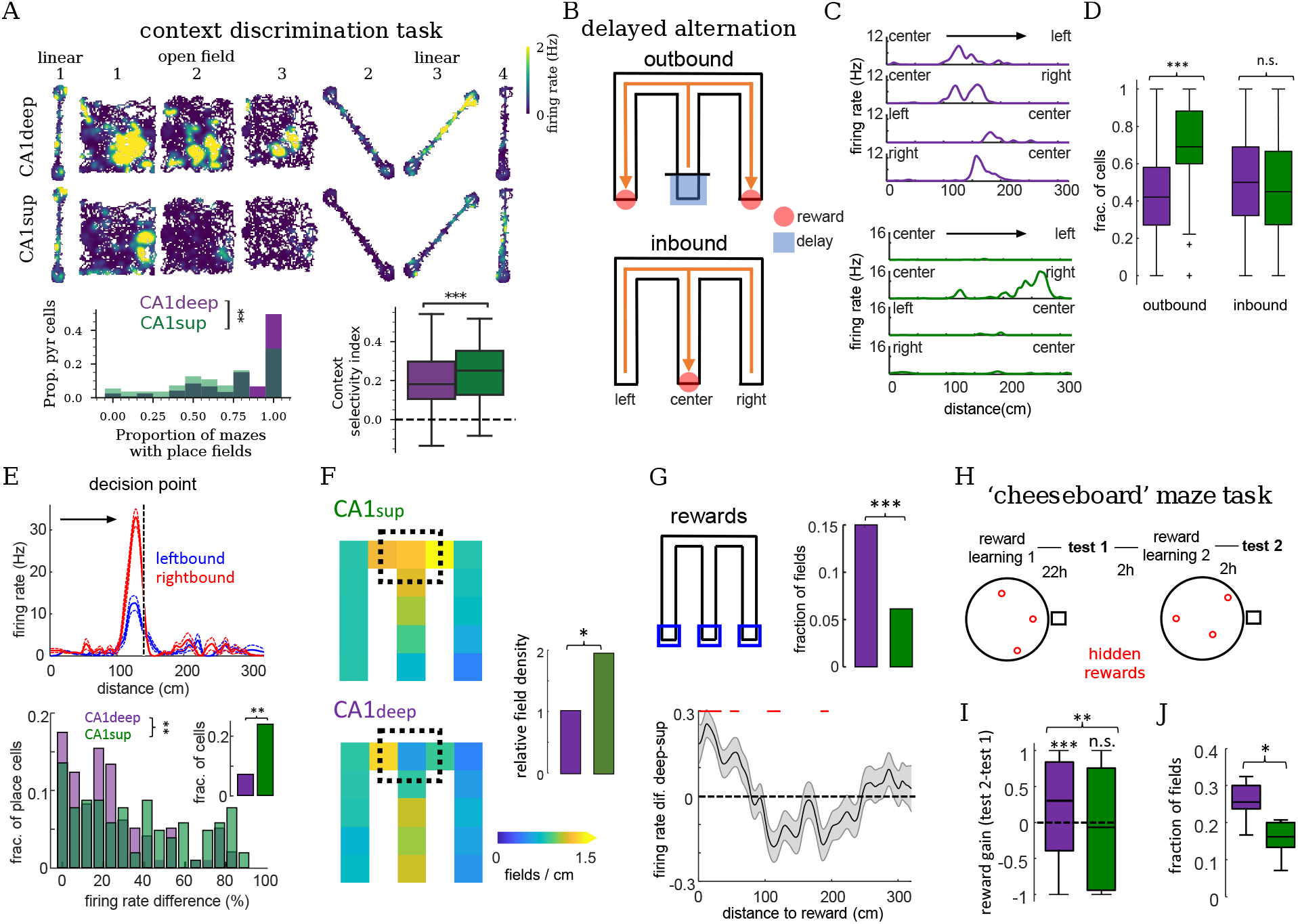
Differential context, choice, and reward coding by CA1deep and CA1sup place cells. **A)** Top: Example rate maps for a CA1deep and CA1sup cell in three different mazes across different contexts. Bottom: fraction of mazes where a cell expressed place fields (*P* = 4.52 × 10^-3^, rank-sum) and context selectivity index (*P* = 8.34 × 10^-4^). **B)** Depiction of delayed alternation task in M-maze. **C)** Example rate maps across M-maze trajectories for a CA1deep (top) and CA1sup (bottom) place cell. **D)** Fraction of place cells with fields in either one of two trajectories in outbound (CA1deep vs CA1sup, *P* = 2.2 × 10^-3^, rank-sum) and inbound (*P* = 0.91) trials. **E)** Top: rate maps for a splitter cell for left and right outbound trials. Bottom: distribution of firing rate difference for left and right outbound trials for cells with overlapping fields in central arm (***n*** = 97 CA1deep / 104 CA1sup; *P* = 0.008) and fraction of splitter cells (inset; *P* = 0.001, Fisher’s test). **F)** Place field density maps for CA1deep and CA1sup place cells in outbound trials (n=398 CA1deep / 422 CA1sup fields). Right: place field density around decision point (black square) / density in rest of maze (*P* = 0.02, Fisher’s test) **G)** Top: Fraction of cells with fields around reward locations (blue squares; n=50 CA1deep / 24 CA1sup fields; *P* = 3.1 × 10^-4^). Bottom: firing rate difference of CA1deep-CA1sup as a function of distance to reward (mean±SEM). Red lines indicate significant differences (*P* < 0.05, t-test with Bonferroni correction). **H)** Depiction of ‘cheeseboard’ maze task. **I)** Increase in firing rate around rewards from test 1 to test 2 (*P* = 4.5 × 10^-5^ & *P* = 0.4 CA1deep/sup sign-rank test, CA1deep vs. CA1sup *P* = 9.6 × 10^-4^, rank-sum). **J)** Fraction of place fields around rewards in test 2 (*P* = 0.015, rank-sum).

To investigate whether memory demands affect the spatial coding properties of CA1deep and CA1sup, we recorded rats on a delayed alternation task on an M-maze (Fig. 3B; n = 6 rats; [18, 40, 41]). In this task, outbound trajectories require animals to remember previous choices and thus are more memory demanding than inbound trajectories. We found that CA1sup place cells (n = 325) tended to be more trajectory-specific than CA1deep place cells (n = 296), which developed fields along multiple trajectories (Fig. 3C and Fig. S7). This difference in trajectory selectivity was only present in the outbound phase, suggesting a memory-dependent phenomenon (Fig. 3D. Trajectory selectivity index CA1deep vs CA1sup: *P* = 8.3 × 10^-6^ outbound/ 0.89 inbound, ranksum). A signature of trajectory-specific coding is the firing rate modulation in a common location of some place cells depending on an animal’s future choice (‘splitter cells’; [40, 42]. To analyze this phenomenon, we focused on cells with place fields in the central arm of the maze. CA1sup place cells exhibited more choice selective firing compared to CA1deep place cells in outbound but not inbound trials (‘prospective splitter cells’; Fig. 3E and Fig. S7). Additionally, we found an overrepresentation of the choice point during outbound trials by CA1sup but not CA1deep place cells (Fig. 3F). Finally, we looked at the representation of reward locations and found that CA1deep cells had a larger proportion of place fields and higher firing rates around reward locations compared to CA1sup (Fig. 3G).

To analyze reward coding, we recorded animals in a “cheeseboard” maze task in which they had to learn new reward locations every day (Fig. 3H; n = 4 rats; [43, 44]). We analyzed the representation of newly learned reward locations by comparing spatial firing in pre- and postlearning ‘test’ sessions without rewards present. CA1deep but not CA1sup place cells selectively increased their firing rates around new reward locations (Fig. 3I) and a larger proportion of CA1deep than CA1sup expressed place fields around new reward locations after learning (Fig. 3J and Fig. S7).

To investigate whether CA1deep and CA1sup preferentially route different task-related information to PFC and MEC, respectively, we detected synchronous assembly patterns across these structures during memory tasks. We found that ~ 11% of the assemblies that emerged during behavior had both hippocampal and cortical members, while the rest were restricted to a single structure (Fig. 4A-B). Hippocampo-cortical assembly members, but not non-member cells, displayed high spike synchrony in a ~ 50ms time window (Fig. 4C). When centered on the activation of hippocampal assembly members, MEC pyramidal cell firing peaked ~ 15 ms later and PFC firing ~ 35ms later, indicating the propagation of information from the hippocampus to the cortex (Fig. 4D and Fig. S8).

**Figure 4:**
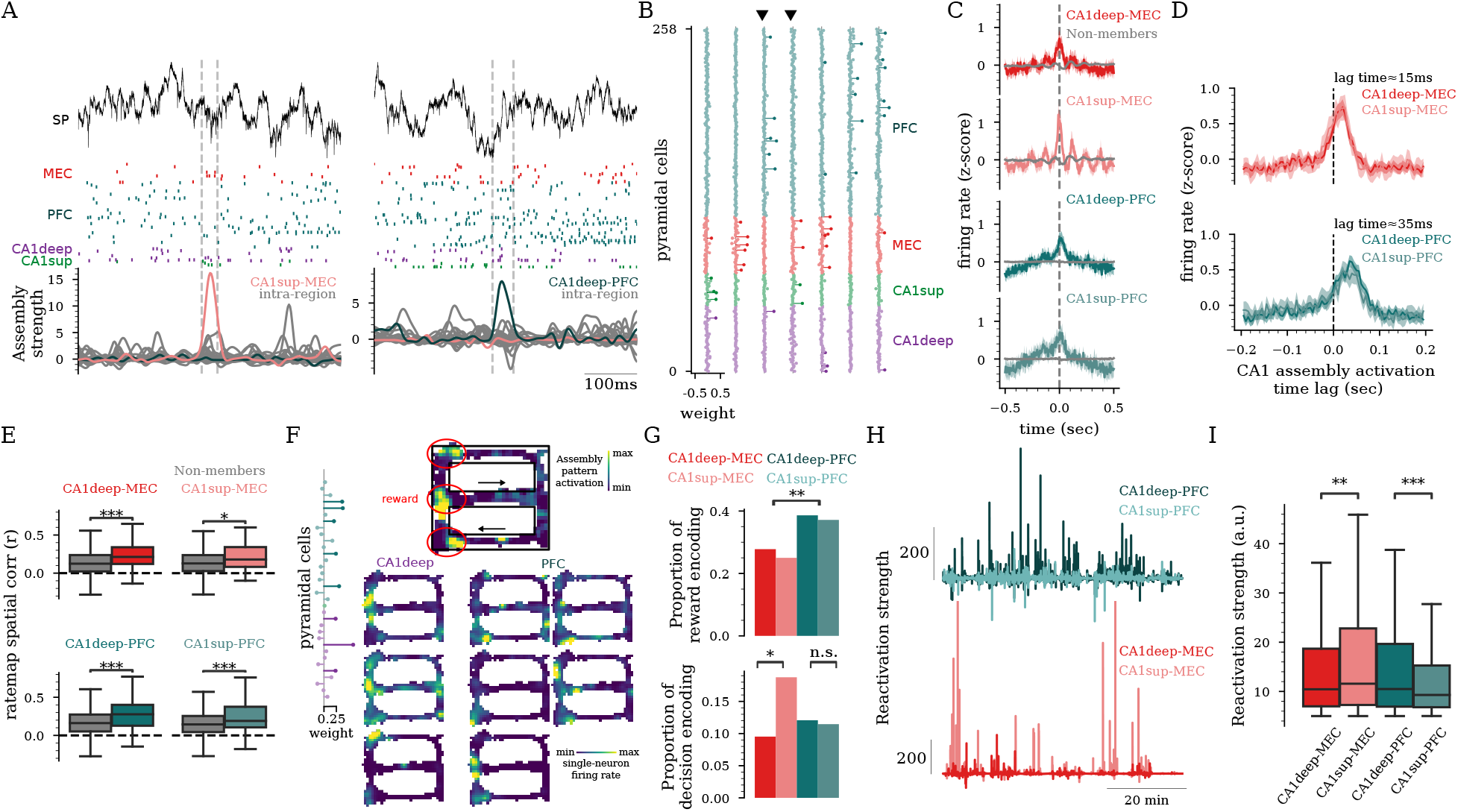
Hippocampo-cortical assemblies encode and reactivate complementary task representations. **A)** Hippocampal LFP (top), unit firing from MEC, PFC, and CA1 (middle), and assembly activation (bottom) during behavior. Dashed lines mark the activation of hippocampo-cortical assemblies. **B)** Example assembly weights including hippocampo-cortical assemblies (marked by black triangles). **C)** Cross-correlograms between spike trains of hippocampo-cortical assembly members and non-members during behavior (Ps < 2 × 10^-16^). **D)** Firing of MEC (top) & PFC (bottom) units center around the discharge of CA1 assembly members (CA1-MEC lag = 15ms; CA1-PFC lag = 35ms). **E)** Rate map spatial correlation for assembly members & non-members (Ps ≤ 1.16 × 10^-2^ rank-sum test). **F)** Example maps for a CA1deep-PFC assembly activation (top) and its individual members firing (bottom) in a delayed alternation task. Red circles indicate reward locations. **G)** Fraction of assemblies encoding reward locations (top; CA1-MEC vs. CA1-PFC *P* = 3.82 × 10^-3^, χ^2^) and decision point in the maze (bottom; CA1deep-MEC vs. CA1sup-MEC: *P* = 0.027, χ^2^). **H)** Example assembly reactivation strength during SWRs. **I)** Reactivation strengths of hippocampo-cortical assemblies in post-task sleep SWRs (CA1deep-MEC vs CA1sup-MEC *P* = 2.49 × 10^-5^; CA1deep-PFC vs CA1sup-PFC *P* = 3.61 × 10^-5^, LME).

We next analyzed the functional correlates of hippocampo-cortical assemblies. Cortical cells had higher spatial firing correlations with hippocampal cells that were members of the same assembly, than with other non-member hippocampal cells (Fig. 4E). Some assemblies were active at all reward locations in the maze while their members fired only at individual reward locations (Fig. 4F), suggesting that cross-regional assemblies can encode generalized representations. Overall, CA1-PFC assemblies displayed higher reward encoding than CA1-MEC assemblies (Fig. 4G). On the other hand, CA1sup-MEC assemblies displayed higher activation around the decision point of the maze than any other assembly type (Fig. 4G).

Finally, we examined the reactivation of cross-regional assemblies during sleep following learning tasks. MEC cells reactivated more strongly with CA1sup than with CA1deep partners while PFC cell’s reactivation was stronger with CA1deep rather than CA1sup (Fig. 4H-I), in line with our previous anatomical and physiological results (Fig. 2).

Experimental and theoretical work has suggested that biological and artificial neural networks composed of diverse elements constitute more efficient memory systems than similar networks integrated by homogeneous units [45, 46]. Hippocampal pyramidal cells exhibit a large molecular, anatomical, and physiological diversity [9–12, 16–20] rooted in development [14, 15]; however, hippocampal memory models do not fully take into account this diversity. Our findings demonstrate that the functional heterogeneity of CA1 pyramidal cells is a major contributor to the flexible computational capabilities of the hippocampus during memory-guided behavior. Contrary to a common tenet that assumes a random cellular organization of hippocampal spatial and memory representations [27–29], we found that pyramidal cells from the same sublayer formed functional assemblies, reactivated together during SWRs and had more similar spatial tuning and functional correlates. CA1deep cells were the main contributors to SWR and assembly dynamics, but CA1sup cells drove the experience-dependent emergence of memory replay, a result that can explain recent observations on the heterogeneous contribution of hippocampal cells to sequence coding [47, 48]. Although previous research showed that SWRs entrain ensemble reactivation in numerous downstream areas [4, 6, 35, 38, 49], until now it was not known if memory representations broadcasted by SWRs are similarly readout in all cortical target regions or if these show selective responses. We reported a previously undescribed anatomical and functional connectivity pattern between CA1, PFC, and MEC. Our results suggest that SWRs carry a multiplexed code in which different types of task-related information are encoded simultaneously but selectively routed to different target areas by deep and superficial pyramidal cells. The existence of parallel hippocampo-cortical subcircuits for selective encoding, routing, and replay of complementary memory representations confers this system with enhanced flexibility and the capability to simultaneously implement different types of computations in support of memory-guided behaviors. These results highlight the importance of incorporating cellular diversity in models of hippocampo-cortical learning and memory functions.

## Acknowledgments

The authors thank Joe Fetcho, Wenbo Tang, Ralitsa Todorova, Laura Berkowitz, Hongyu Chang, Jesse Goldberg, Lucas Sjulson, and Nilay Yapici for useful comments on the manuscript and Melisa Warden for providing access to the confocal microscope. The authors also thank Roman Huzsar and Yunchang Zhang for advice on histological quantification, and Mazen Kheirbek for advice on retrograde tracing experiments. Funding: This work was supported by NIH grant 4R00MH120343, Sloan Fellowship, Whitehall Research Grant and Klingenstein-Simons Fellowship (AFR), NIH grant 4R00MH122582, BBRF NARSAD Award (AO), NYSTEM C029155, and NIH S10OD018516 (Cornell University BRC Imaging Facility).

## Author contributions

A.F.-R., A.O., and R.E.H. conceived and designed experiments and analyses; R.E.H., H.L.R., C.L., A.O., and A.F.-R. performed experiments and analyzed data; A.F.-R., A.O., and R.E.H. wrote the manuscript.

## Materials and methods

### Surgical procedures

Rats (adult male Long-Evans, 300-500 g, 3-6 months old) were kept in the vivarium on a 12-hour light/ dark cycle and were housed 2 per cage before surgery and individually after it. All experiments conformed to guidelines established by the National Institutes of Health and have been approved by the Cornell University Institutional Animal Care and Use Committee.

Silicon probe implantation were performed as described previously [18, 44]. Animals were anesthetized with isoflurane anesthesia and craniotomies were carried out under stereotaxic guidance. Silicon probes (NeuroNexus, Cambridge Neurotech, or Diagnostic Biosignals) were mounted on custom-made 3D-printed micro-drives to allow precise adjustment of the vertical position of sites after implantation. The probes were inserted above the target region. Craniotomies were sealed with sterile wax. Two stainless steel screws were placed bilaterally over the cerebellum to serve as ground and reference electrodes. Several additional screws were driven into the skull and covered with dental cement to strengthen the implant. Finally, a copper mesh mounted on a 3D-printed resin base was attached to the skull with dental cement and connected to the ground screw to act as a Faraday cage, attenuating the contamination of the recordings by environmental electric noise and protecting the headgear. After post-surgery recovery, probes were moved gradually in 50 to 150 *μ*m steps per day until the desired position was reached. Hippocampal, prefrontal cortex, and entorhinal cellular layers were identified physiologically by unit activity and characteristic LFP patterns [21, 50].

A variety of different silicon probes were implanted in the dorsal hippocampus (−4.5-4.0mm antero-posterior from Bregma - AP and 2.6mm from midline; ML), the medial entorhinal cortex (−9.0mm AP, 4.0mm ML), and prefrontal cortex (3.0mm AP, 0.7mm ML). Data from some of the animals included in this study have also been included in previous studies, and surgical procedures have been described in more detail there [18, 21, 35, 44, 47, 51–56].

### Optogenetic experiments

The optogenetic experiments analyzed here were performed as part of a previously published study by Fernandez-Ruiz et al., [18]. Briefly, rats (n = 5) were injected with AAV5-CaMKIIa-hChR2(H134R)-EYFP from UNC Vector Core (a gift from Dr. Karl Deisseroth). Three injections of 150 nL each were performed along the longitudinal axis of both dorsal hippocampi, right above the CA1 pyramidal layer. After injection, craniotomies were sealed and animals recovered in the vivarium for three weeks. Following this period, a second surgical procedure for implanting optic fibers and electrodes was performed. 200 *μ*m core multi-mode optic fibers (Thor Labs) were implanted in the same craniotomies used previously for virus injection, right above the CA1 pyramidal layer. Optic fibers were directly coupled to 460 nm blue light-emitting laser diodes (PL-450, Osram). A 64-channel silicon probe was also implanted targeting the CA1 region.

A closed-loop system was used in order to detect SWR during behavior and trigger light stimulation. Once a SWR was detected, a small amplitude, 100-ms, trapezoidal light pulse was delivered simultaneously through all fibers. The intensity was manually adjusted in each animal during a test session in the home cage by gradually increasing light power until a clear ripple was evoked. Pulses were delivered every time a SWR was detected while the animal was performing the task in the M-maze (see below).

### Behavioral recordings

After surgery, the animals were handled daily and accommodated to the experimenter, recording room, and cables for one week before the start of the experiments. Prior to the start of the behavioral experiment, the animals were water restricted. Physiological signals were recorded during five different tasks.

In the linear or circular track, rats (n = 27) were trained to run back and forth to collect a small water reward. After the first one or two days, all animals performed 40-80 trials per day. The session was terminated when the animal was satiated, typically for one hour. The linear track was placed one meter above the floor and was 240-cm long and 7-cm wide with 5-10cm walls. The circular track was 100 cm in diameter. Water rewards were delivered in a predetermined position only when the animals had performed a full clockwise run.

In the M-maze experiment, rats (n = 7) were first pre-trained to collect sugar-water rewards in the linear track. Once they were used to running and obtaining rewards, the M-maze recordings started [18]. In the M-maze, rats were rewarded with sugar water each time they reached the end of the three arms in the correct task sequence (center-left-center-right-center). The two components of the task (outbound and inbound) were evaluated separately to assess task performance [18]. The animal was confined at the end of the central arm for 20 s after each inbound trial. Animals performed the task for 10 days. Two 30-minute sessions were conducted in the morning and afternoon, separated by 5 h and preceded and followed by ~ 1 h sleep recording sessions.

Cheeseboard Maze (120 cm diameter), where the animals (n = 9) learned to find 3 goal wells that contained water rewards. A trial was completed once the animal had retrieved all rewards and returned to the start box to collect an additional food pellet reward. The locations of the goal wells changed daily but were fixed within a session. This strategy required the animals to update their memory for the new goal locations in each session but in an otherwise familiar environment. Note that between trials, there was always a delay of approximately 30 seconds. A pre-probe session (‘test 1’), without rewards present, was run every day to assess whether the animal remembered the previous day’s positions. Following the learning session, a post-probe session (‘test 2’), also without rewards, was also conducted to examine whether the animal remembered the newly learned locations. The inclusion of data for the analysis required that the animal be pre-trained for a week. In addition, the rat’s performance in each included session had to show a trend of learning across trials and memory performance in probe sessions was above chance levels. Sleep sessions were recorded before and after each learning session.

In the rectangular figure-8 maze (or ‘T-maze’) animals (n = 15) were trained to alternate between right and left arms to collect water rewards. At the beginning of the trial, rats were confined in the start box for 15 s. After opening the door that led to the central arm, the animals ran through the central arm and chose to turn right or left. If the choice was correct (opposite arm than visited in the previous trial), they would find a water reward at the end of the side arm; if incorrect, the reward port would remain empty. Only sessions when animals had been trained in the task for at least one week and performed above 70% correct trials were included. The maze was placed one meter above the floor and had 160 cm in length (for the central and side arms) and 134 cm in width (for the lateral connecting arms).

Open field environment (180 cm × 180 cm, or 120 cm × 120 cm), where the animals (n=10) randomly foraged for pieces of randomly placed sugar pellets until they fully sampled the entire environment (~30 minutes).

Plus maze environment (100 cm × 100 cm), where the animals (n=1) in which the rats were motivated to run to the ends of four corridors, where water was given every 30 s.

Novelty sessions took place in a different room never visited before by the animal. The ‘familiar’ and ‘novel’ rooms had unique prominent distal cues. Different behavioral apparatuses (described above) were used during novelty sessions: linear and circular tracks, open field, T-maze, and cheeseboard maze. Animals were trained to run in these mazes to collect water or food rewards until they were satiated, typically 30-60 minutes. All animals were trained for multiple days in the familiar room and behavioral apparatus, which differed from those used in novelty sessions for each animal.

For all mazes, the position of the animal was recorded with an overhead camera (Basler) and tracked with DeepLabCut [57] or custom software.

In the datasets obtained from the CRCNS repository (http://crcns.org/data-sets/hc/hc-3, http://crcns.org/data-sets/pfc/pfc-2, https://crcns.org/data-sets/hc/hc-11, https://crcns.org/data-sets/hc/hc-14), animals were also recorded during linear track, open field, and T-maze alternation task that have been described in detail previously [35, 47, 54–56, 58–60].

### Tissue processing and immunohistochemistry

Following the termination of the experiments, animals were deeply anesthetized and perfused transcardially first with 0.9% saline solution followed by 4% paraformaldehyde solution. The brains were sectioned into 70-*μ*m thick slices (Leica Vibratome). The sections were washed and mounted on glass slides with a fluorescence medium (Fluoroshield with DAPI - F6057, Sigma, USA). A confocal microscope (Zeiss LSM 800) was used to obtain high-quality photos.

### Retrograde tracing experiments

For CTB retrograde studies, 330nl of conjugated CTB (Life Technologies, Carlsbad, CA) was injected into the prefrontal cortex (PFC, CTB Alex Fluor 488) and the Medial Entorhinal Cortex (MEC, CTB Alex Fluor 555) in the same animal in one surgery [61]. For each site, there were 10 injections of 33nl at 23 nl/sec with a 15-sec delay. Injection sites (in mm from bregma) for PFC were AP +3.0, ML +1.0, DV −1.5, −1.8, −2.0, −2.3, −2.7, −3.1, −3.6, −4.1, −4.5, −5,0. Injection sites (in mm from bregma) for MEC (1) AP −7.7, ML + 4.6, DV −4.7; (2) AP −8.0, ML +4.6 DV −4.3, −4.8, −5.3; (3) AP −9.1, ML +4.6, DV −3.2, −3.7, −4.2, −4.5, −4.7, −5.2. Animals were perfused transcardially 7 days after injection, first with 0.9% saline followed by 4% paraformaldehyde solution. The brains were sectioned in 70-*μ*m thick slices (Leica Vibratome), then washed with DAPI (Thermo Fisher Scientific), and mounted on glass slides. Images were taken on Confocal microscopes (Zeiss LSM 800 for quantification; Zeiss LSM880 i880 for high-resolution images(Cornell University Biotechnology Resrouce Center Imaging Core)). Injection sites were verified for both PFC and MEC (Fig.2A).

Quantification was performed first in ImageJ by taking CA1 segments from slices AP −3.2 to −4.5 mm and rotating the layer horizontally (Fig. S4). The border between the stratum radiuatm (SR) and the pyramidal layer (SP) and the border between the stratum oriens (SO) and the SP were then marked according to the DAPI staining marking in alignment with the Mouse Brain Atlas [62] (Fig. S4). The distance between SR-SP and SO-SP was taken as the pyramidal layer thickness. The distance of each cell (Alex Fluor 488-PFC+ and Alex Fluor 555 MEC+) from the SR-SP. border was divided by the pyramidal layer thickness to calculate the normalized depth of all labeled cells within the CA1 pyramidal cell layer [15].

### Recording and data processing

Recordings were conducted using the Intan RHD2000 interface board or Intan Recording Controller, sampled at 20 kHz. Amplification and digitization were done on the head stage. Data were visualized with Neurosuite software, Neuroscope. All local field potential (LFP) analyses (SWR detection, state scoring, etc.) were conducted on the 1,250-Hz down-sampled signal.

## Quantification and statistical analysis

### Spike sorting and unit classification

Spike sorting was performed semi-automatically with KiloSort [63] (https://github.com/cortex-lab/KiloSort), followed by manual curation using the software Phy (github.com/kwikteam/phy) and custom designed plugins (https://github.com/petersenpeter/phy-plugins) to obtain well-isolated single units. Cluster quality was assessed by manual inspection of waveforms and auto-correlograms, and by the isolation distance metric. Multiunit, noise clusters or poorly isolated units were discarded for analysis. Well isolated units were classified into putative cell types using the Matlab package Cell Explorer (petersenpeter.github.io/Cell-Explorer [64]). Spiking characteristics, including the autocorrelograms, spike waveforms, and putative monosynaptic connections derived from short-term cross correlograms, were used to select and characterize well-isolated units. Three cell types were assigned: putative pyramidal cells, narrow, and wide waveform interneurons. Two key metrics used for this separation were burst index and trough-to-peak latency (Fig. S1). Burst index was determined by calculating the average number of spikes in the 3–5 ms bins of the spike autocorrelogram divided by the average number of spikes in the 200–300 ms bins. To calculate the trough-to-peak latency, the average waveforms were taken from the recording site with the maximum amplitude for the averaged waveforms of a given unit.

### Classification of deep and superficial CA1 pyramidal cells

To classify all recorded CA1 pyramidal cells based on the radial position of their soma, the relative depth of each electrode across the pyramidal layer was first determined. This classification was based on the stereotypical depth profile of SWRs as the probe transects the pyramidal layer. The slow envelope of the SWR has a positive polarity above the pyramidal layer a reverts that polarity right across the middle of the layer [17, 21]. Below the middle of the pyramidal layer (superficial sublayer), SWR envelope has a negative polarity. The amplitude of this envelope gradually increases, peaking in the middle of the stratum radiatum (the sharp-wave). The relative depth of each electrode was determined in relation to the channel of polarity reversal (the middle of the layer: 0 *μ*m), considering the know inter-electrode distance. The classification assigned three labels: deep, middle, and superficial defined as deep: < −30*μm*, middle: > −30*μm*& < 30*μm*, superficial: > 30*μm*. The position of the soma of each pyramidal cell was determined as the electrode with the largest amplitude waveforms across the shank.

### SWR detection

To detect SWRs, the wide-band signal was band-pass filtered (difference-of-Gaussians; zero-lag, linear phase FIR), and instantaneous power was calculated by clipping at 4 SD, rectified, and low-pass filtered [50]. The low-pass filter cut-off was at a frequency corresponding to *p* cycles of the mean bandpass (for 80-250 Hz band-pass, the low-pass was 55 Hz). Subsequently, the power of the non-clipped signal was computed, and all events exceeding 4 SD from the mean were detected. The events were then expanded until the (non-clipped) power fell below 1 SD; short events (< 15ms) were discarded. Sharp waves were detected separately using LFP from a CA1 str. radiatum channel, filtered with band-pass filter boundaries (5-40 Hz). LFP events of a minimum duration of 20 ms and a maximum of 400 ms exceeding 2.5 SD of the background signal were included as candidate SPWs. Only if a sharp wave was simultaneously detected with a ripple was a CA1 SPW-R event retained for further analysis.

### SWR spike content analysis

Well-isolated putative units with at least 100 spikes in a given session were included in the analysis. Within-SWR firing rate was calculated as the number of spikes within SWRs divided by the cumulative SWR duration for the session. The probability of participation of individual units in SWRs was defined as the number of events in which a neuron fired at least one spike during the SWR divided by the total number of SWR detected. Cross-correlograms (CCG) were constructed with within-SWR spikes as a function of the latency binning them into 5 ms size bins. Spike counts in CCGs of all pairs in each category were z-scored. For peri-SWR firing histograms, single unit firing rates were z-scored, averaged, and plotted with ± 95% confidence intervals.

### Place cell analysis

Spiking data and the tracked animal’s position were binned into 3-cm wide segments of the camera field projected onto the maze floor, generating raw maps of spike counts and occupancy. A Gaussian kernel (SD = 3 cm) was applied to both raw maps of spike and occupancy, and a smoothed rate map was constructed by dividing the spike map by the occupancy map. Independent rate maps were constructed for the different running directions in the mazes. Only periods in which the animal velocity was greater than 4 cm/s were included. A place field was defined as a continuous region of at least 15cm^2^, where the mean firing rate was above 10% of the peak rate in the maze, and the peak firing rate was > *3Hz*. We calculated the spatial information encoded in place cell firing rates using the method of [65] to calculate spatial information in bits per spike as follows:

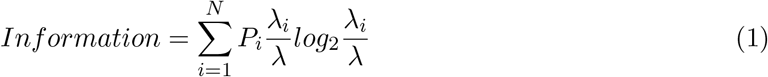

Where the environment is divided into spatial bins *i* = 1,…, *N, P_i_* is the occupancy probability of bin *i, λ_i_* is the mean firing rate for bin *i*, and *λ* is the overall mean firing rate of the cell.

In the delayed alternation task, a ‘trajectory selectivity index’ (TSI) was calculated similarly to previous reports (e.g. [4]). First, rate maps were constructed separately for left and right bound trials. Then the firing rate difference between different trajectories was compared (i.e., center-to-left versus center-to-right outbound trajectories, and left-to-center versus right-to-center inbound trajectories) as:

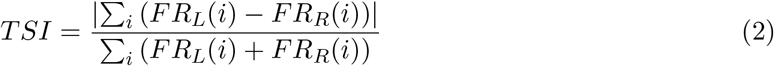

where *FR_L_*(*i*) is the firing rate in the *i*-th spatial bin on the stem during left trials (i.e., center-to-left trial for outbound, or left-to-center trial for inbound), and *FR_R_*(*i*) is for the right trials.

To identify ‘splitter cells’ place cells with fields in the central arm of the maze were selected. If the cell had overlapping place fields in both directions with an average peak firing rate difference of at least 50% it was considered a splitter cell.

For reward location of decision point-related analysis, place fields were considered if their peak fall within 20cm of those landmarks. ‘Reward gain’ in the cheeseboard maze was calculated as the firing rate of a cell in a 20cm radius around previously rewarded locations divided by the firing in the rest of the maze.

### Cell assembly analyses

Cell assemblies were identified as previously described [32, 44, 66–69]. Significant co-firing patterns were detected using an unsupervised statistical method based on independent component analysis (ICA). The spike trains for each neuron were binned into time windows (25-ms for inter-hippocampal in Fig. 1 and 50-ms for inter-region in Fig. 4) and *z*-score transformed to eliminate biases due to differences in average firing rates. Next, a principal component analysis was applied to the binned spike matrix (*Z*). The correlation matrix of *Z* was given by 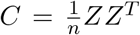 and the eigenvalue decomposition of C was given by:

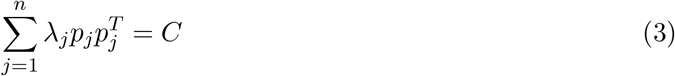

where *λ_j_* is the eigenvalue of *C* and *p_j_* is its corresponding eigenvector. The Marcenko-Pastur law was used to estimate the number of significant patterns embedded within Z. For a *nXB* matrix, an eigenvalue exceeding *λ_max_*, defined by 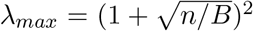, signifies that the pattern given by the corresponding principal component explains more correlation than would be expected if the neurons were independent of each other [70]. The number of eigenvalues exceeding λ_mαx_ was defined as *N_A_* and therefore represents the minimum number of distinct significant patterns in the data [6, 66]. The significant principal components were then projected back onto the binned spike data

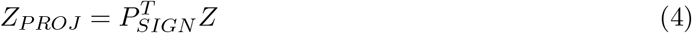

where *P_SIGN_* is the *nXN_A_* matrix with the *N_A_* principal components as columns.

Independent component analysis (ICA), using the fast ICA algorithm [71], was then applied to the matrix *Z_PROJ_*. That is, an *N_A_XN_A_* unmixing matrix *W* was found such that the rows of the matrix *Y* = *W^T^Z_PROJ_* were as independent as possible. The unmixing matrix W was then used to derive each cell’s weight within each assembly *V* = *P_SIGN_W*.

Cell assembly members were identified using Otsu’s method [68, 72] to divide the absolute weights into two groups that maximized inter-class variance. Neurons in the group with greater absolute weights were classified as members. The goodness of separation was quantified using Otsu’s effectiveness metric, namely the ratio of the inter-class variance to the total variance. This procedure yielded a set of vectors *C_i_* representing the detected cell assemblies.

To determine the strength of the expressed assemblies, we tracked each assembly pattern *v_k_* over time by:

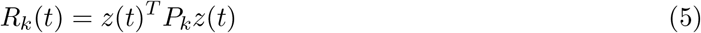

where *z*(*t*) is a smooth vector-function containing for each neuron its *z*-scored instantaneous firing-rate and *P_k_* is the matrix projecting *z*(*t*) to the activation-strength of the assembly pattern *k* at time t. To increase the temporal resolution, *z*(*t*) was obtained by convolving the spiketrain of each neuron with a Gaussian kernel 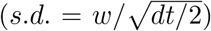 and then *z*-scoring each trace. *w* was set to 25 ms to match the bin size used to identify the assembly patterns. Assembly-activations were defined as peaks exceeding *R_THRES_* > 5.

The within SWR assembly reactivation strength was defined as the median peak activation value of events with R *>* 5 that occurred during SWR epochs in post-task sleep. Only sessions with ≥ 25 SWR events were considered for assembly analysis to assess distribution parameters.

To determine the spatial encoding of assemblies, time-resolved assembly activations were aggregated over spatial 3-cm bins which created 2-D assembly maps (Fig. 4F).

### Detection of brain states

State scoring was performed as previously described in Levenstein et al., [73]. Briefly, the local field potential (LFP) was extracted from wide-band data by lowpass filtering (sinc filter with a 450 Hz cut-off band) and downsampling to 1250Hz. Broadband LFP, narrow-band theta frequency LFP, and estimated electromyogram (EMG) were used for state scoring. Spectrograms were computed from broadband LFP with a Fast Fourier transform in 10s sliding windows (at 1s), and a principle component analysis was computed after a Z-transform. The first principle component reflected power in the low (<20Hz) frequency range, with oppositely weighted power at higher (>32Hz) frequencies. Theta dominance was quantified as the ratio of powers in the 5–10Hz and 2-16Hz frequency bands. EMG was estimated as the zero-lag correlation between 300-600Hz filtered signals between recording sites [74]. Soft sticky thresholds on these metrics were used to identify states. High LFP principal component 1 and the low EMG were considered NREM, the high theta and low EMG were considered REM, and the remaining data were taken to reflect the waking state.

### Detection of replay

A Bayesian decoding approach (see [75] for the original method) was used to detect and analyze replay events (see [47, 76–80] for the use cases of this method to investigate replay). Spike counts from each SWR were first binned into 20ms time bin t from *N* units *O_t_* = (*o*_1*t*_*o*_2*t*_…*o_Nt_*) (200ms time bins were used for active behavior decoding). Then the posterior probability distributions for each t over binned positions (3 cm) along the linear track were calculated using Bayes’ rule:

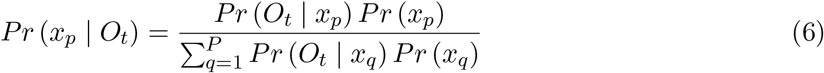

Where *x_p_* is the center of the *p – th* position bin. Because we assumed Poisson firing, the prior probability, *Pr*(*O_t_|x_p_*,), for the firing of each unit *n* is equal to

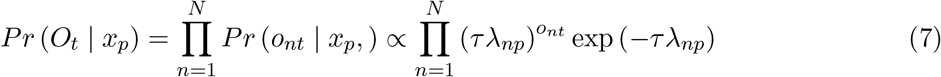

where *τ* is the duration of time (20 ms) and *λ_np_* is the mean firing rate of the *n – th* unit in the *p – th* position. We assumed a uniform prior distribution *Pr*(*x_p_*) over the position bins. This implementation of Bayesian decoding was carried out using tools in the Python package Nelpy [81].

Each replay candidate was considered a significant replay event if the series of decoded positions were more consistent with an ordered trajectory compared to a surrogate distribution. Linear regression was used to fit a line to the posterior probability distribution. A Bayesian replay score for a given event was defined as the sum probability mass under the fit line within a bandwidth (21 cm) [77]. For each candidate event, we generated 1500 surrogates of the posterior probability distribution by circularly shifting each column of the posterior probability matrix by a random amount. A Monte Carlo p-value for each event was obtained from the number of surrogate events with replay scores higher than the observed score. The threshold for significance was held at 0.05. Replay events were identified from events with at least 5 active pyramidal cells with a peak firing rate of ≥ 1Hz and a peak-to-mean firing rate ratio of 1.5. In addition, candidate events with many (> 50%) inactive spike count bins were discarded.

This Bayesian decoding algorithm was also used to estimate the animal’s location during active running (speed > 4 cm/s), similar to previous studies [77, 82–84].

Recording sessions with poor decoding accuracy of the animal’s position were excluded from the replay analysis. Decoding accuracy was assessed with a surrogate analysis, where the spike times of each unit were circularly shifted by a random amount. This was done to preserve the temporal components of each spike-train, but to randomize the spike times relative to the position in which they fired. Linear regressions between the real and decoded position of the animals were calculated, and a Monte Carlo p-value for each recording session was obtained using the R^2^ values of each regression. Only sessions with a significant Monte Carlo p-value < 0.05 were used to quantify replay.

### Predicting downstream neural activity

To predict neural activity from cortical regions downstream of CA1, we used kernel reduced rank regression (Fig.2K-L, [85–89]). This is a form of regularized linear regression, with the prediction weights matrix restricted to a specific rank, reducing the number of parameters and making it more robust to overfitting. For analyses where cortical activity was predicted from CA1 during SWRs, spike counts were binned −50ms to +200ms from the start and stop of the SWR respectively to allow for synaptic lag times (Fig. 2F and Fig. 4D). A 5-fold cross-validated grid search on held-out data (60% training data, 40% testing data) was used to locate the optimal rank parameter per recording session using GridSearchCV from scikit-learn [71].

To predict the relative proportion of CA1deep and CA1sup neurons that were active in each SWR, cross-validated ridge regression (RidgeCV from scikit-learn [71]) was used for its robustness to highly collinear predictors. Briefly, a binary spike matrix *X* (n-units by n SWRs) was constructed from cortical (PFC or MEC) activity ± 100ms around each SWR. Dimensionality reduction was then applied to X using PCA to reduce the explained variance of X to 80% in order to control for different numbers of units from session to session and between cortical regions. From here, the resulting matrix was split into testing and training sets (40/60 split) and used to predict the relative proportion of CA1deep and CA1sup neurons that were active in each SWR (y). The accuracy of this prediction was then compared to the accuracy from a shuffled model where the SWR identify was randomly permuted 1000 times resulting in a prediction gain metric (Fig.2G).

Another approach was used to predict the relative proportion of CA1deep and CA1sup neurons that were active in each SWR. The slopes from a linear regression that modeled the relationship between the relative proportion of CA1deep and CA1sup neurons and the number of active cortical neurons in each SWR were compared between MEC and PFC (Fig. 2H). Similarly, the slopes from a regression that modeled the relationship between the relative proportion of CA1deep and CA1sup neurons and the average firing rate of cortical neurons in each SWR were compared between MEC and PFC (Fig. 2I). For both metrics, if the number of CA1deep neurons was positively related to downstream firing, slopes would be positive. Likewise, if the number of CA1sup neurons was positively related to downstream firing, slopes would be negative.

Mutual information was used to measure the relation between CA1deep/CA1sup and cortical activity. Briefly, firing rates within SWRs from CA1deep, CA1sup, MEC, and PFC pyramidal cells were calculated and pairwise mutual information was calculated between the resulting vectors. Time-series mutual information between the two vectors *x* and *y* was calculated according to Cover & Thomas [90] as:

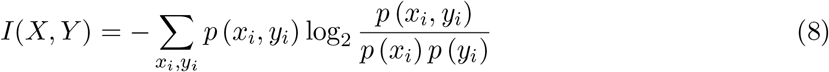

### Theta-cycle detection

As theta phase shifts along the radial CA1 axis [91], a channel located above the center of the pyramidal layer was selected for consistency of the extracted phases across recordings. Theta cycles were calculated similar to previous studies [92]. To the broadband LFP, downsampled at 1250 Hz, we applied a broad bandpass filter (1–80 Hz) and determined local minima and maxima in the theta range. Local minima and maxima were used to determine the duration of the ascending and descending parts of the theta cycle to calculate the asymmetry index. Cycles below 100 ms and above 200 ms were discarded. From this, local minima and maxima were determined to extract the waveform phase by linear interpolation between them, with minima equaling 0° and maxima equaling 180°. With this approach, regardless of the length of the ascending and descending parts of the theta cycle, each part always covers 180°.

### Co-firing analysis

The spike count was calculated in each interval (either a SWR or a theta cycle), resulting in a vector of spike counts for each pyramidal neuron. Pearson’s correlation coefficients between spike count vectors of different neurons were calculated to estimate the SWR and theta-cycle co-firing of each pair.

### Context-selectivity index

To estimate how well individual neurons were able to differentiate different environments, a classifier model (ExtraTreeClassifier in scikit-learn [71, 93]) was used. Briefly, binned spike-trains (dt = 30s, smoothing sigma = 15s) were used to predict which environment they were sampled for recording sessions with at least 2 environments. The accuracy obtained from the test set (33% of the data) was then compared to the chance levels (Fig. 3A). For example, if 70% of bins were accurately predicted and there were 2 environments for the recording session, giving a chance level of 50%, the context selectivity index was 0.2.

### Statistical analysis

Statistics were calculated using R, Python, and Matlab. Using R packages lme4 [94] and lmerTest [95], linear mixed-effects models were used to assess group differences as they account for correlated data, such as many neurons and sessions from the same rat when conducting statistical modeling and hypothesis testing [96]. The basic model used was *feature ~ group + (1|animal/session)*. The normality and independence of the errors were visually inspected and corrected with an appropriate transform if the errors were skewed. Model assumptions were also inspected using the R package DHARMa [97]. When appropriate, a rank-sum test was used. No data were excluded based on statistical tests. Group differences in proportion were modeled using generalized linear models with mixed effects and fit using the glmer() function with family = ‘‘binomial”. When appropriate, a Chi-square (χ^2^) or Fisher’s exact test was used. The significance threshold was set at α = 0.05 unless otherwise noted.

In box plots, the central mark indicates the median; the bottom and top edges of the box indicate the 25th and 75th percentiles,

No specific analysis to estimate minimal population sample or group size was used, but the number of animals, sessions, recorded cells, and SWR events were larger or similar to those employed in previous related works [4, 6, 17, 18, 24, 25, 33, 44, 49]. The unit of analysis was typically identified as single cells or SWR events. In some cases, the unit of analysis was sessions or animals, and this is stated in the text.

**Figure S1:**
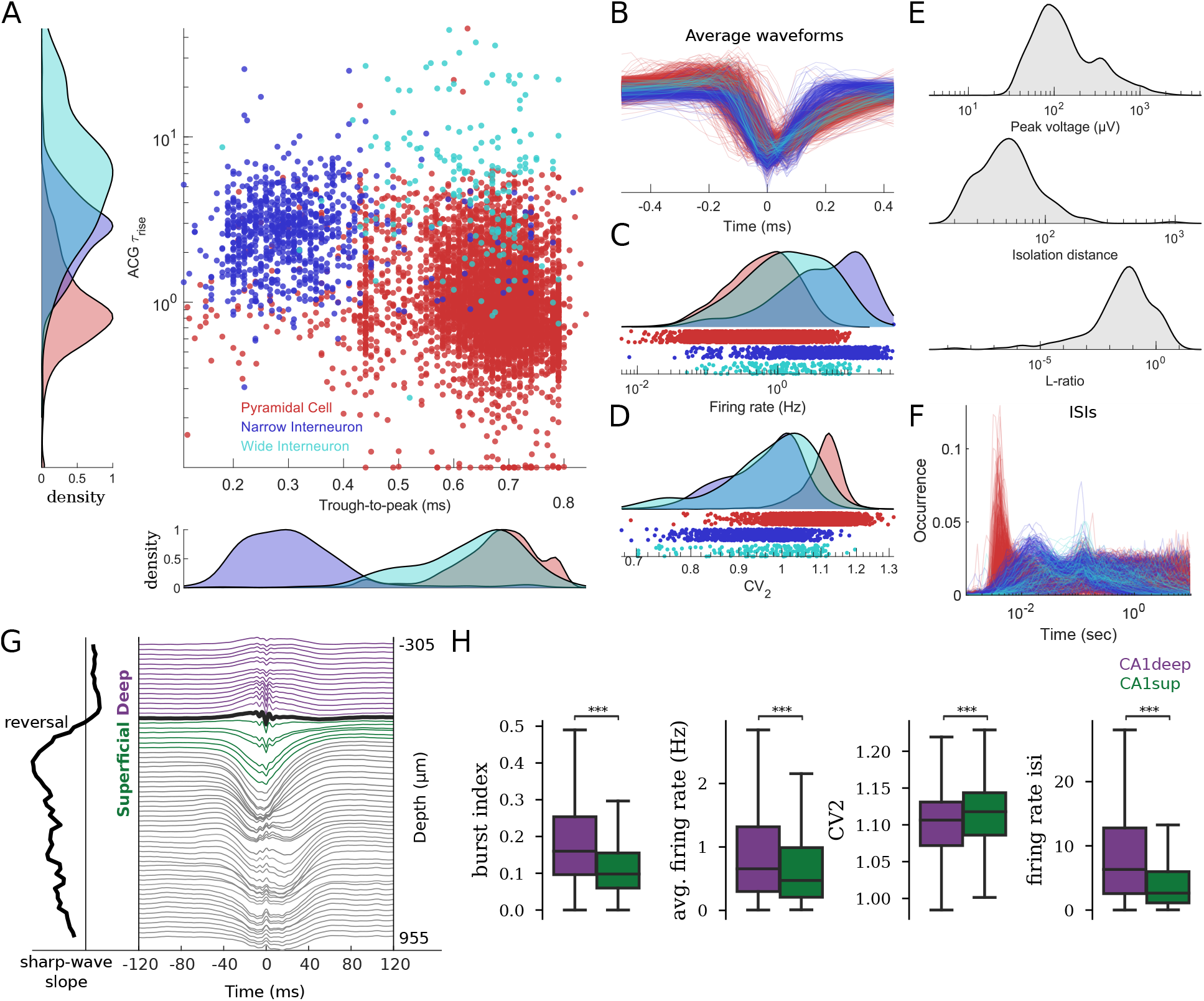
Pyramidal cell versus interneuron classification and intrinsic firing properties. **A)** Average waveform trough-to-peak duration versus autocorrelogram burst metric (ACG *τ_rise_*) of putative pyramidal cells (n=5,245), narrow interneurons (n=823), and wide interneurons (n=171) from 27 rats. **B-F)** Intrinsic physiological metrics for pyramidal cells and interneurons. **B)** Average waveforms. **C)** Average firing rate. **D)** Coefficient of variation 2 (CV2) [98]. **E)** Peak voltage of average waveform, unit isolation distance, and L-ratio. **F)** Inter-spike-intervals. **G)** Example SWR depth profile and CA1 deep/superficial channel classification from a linear probe. **H)** intrinsic firing properties for CA1deep (n=3,658) and CA1sup (n=1,946)); Burst index [59] (*P* < 2 × 10^-16^, LME), average firing rate (*P* < 2 × 10^-16^), CV2 (*P* = 2.21 × 10^-8^), and inter-spike interval (*P* < 2 × 10^-16^).

**Figure S2:**
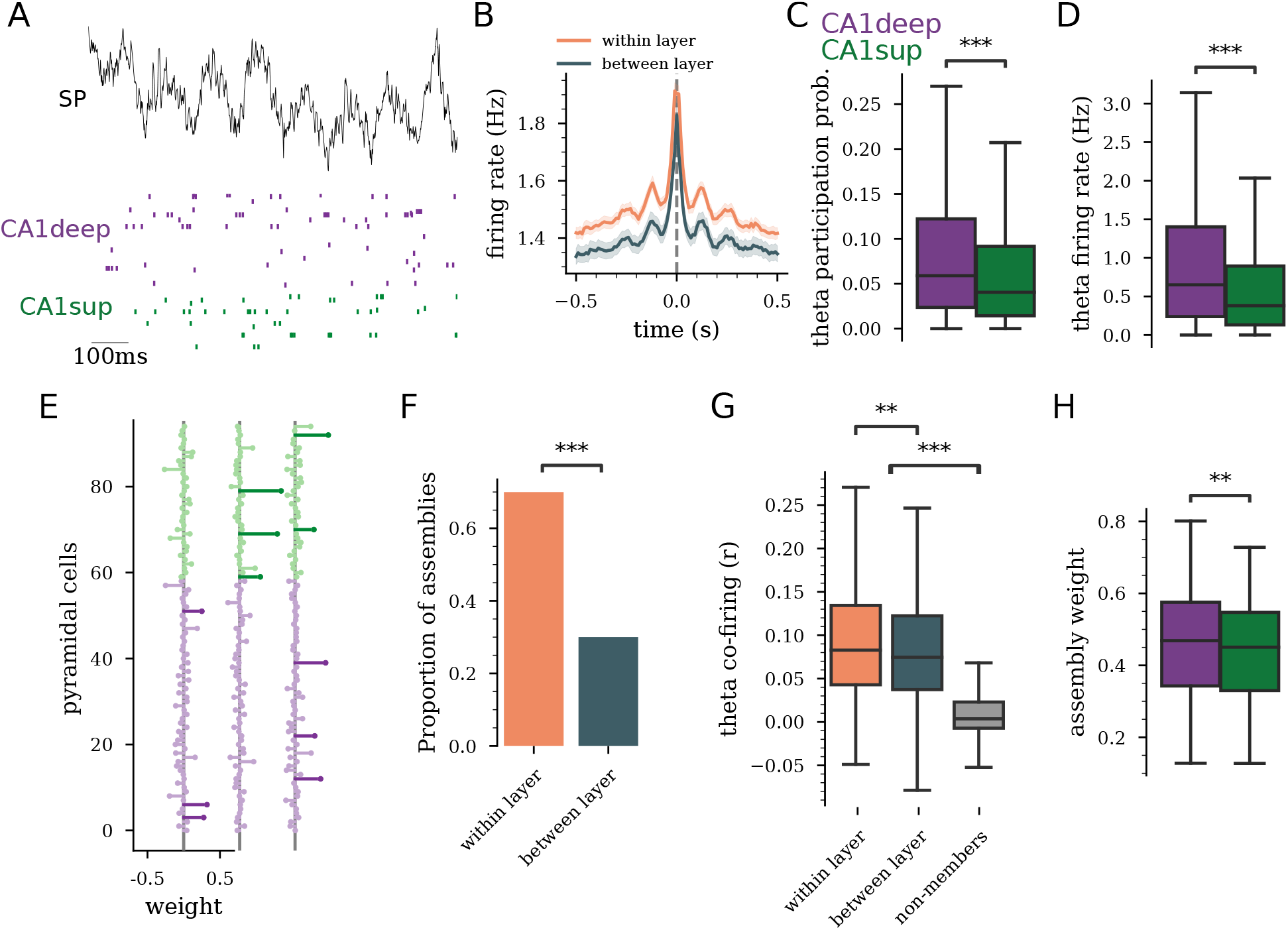
CA1deep and CA1sup co-firing in theta epochs. **A)** Example LFP trace (top) during theta epoch and CA1deep/CA1sup raster (bottom). **B)** Spike-train cross-correlogram during theta epochs for all pyramidal cells (mean±95%CI; n=87,796 pairs from 33 rats). **C)** Participation probability in theta cycles (*P* < 2 × 10^-16^). **D)** Firing rate in theta cycles (*P* < 2 × 10^-16^). **E)** Example assembly weights. The length of horizontal lines indicates the contribution of each cell. Significant members are darker. **F)** Assembly member composition (n = 946 assemblies, *P* = 1.16 × 10^-67^, χ^2^). **G)** Within and across layer theta cycle co-firing (*P* = 1.53 × 10^-3^, rank-sum test) of assembly members and non-members (*P* < 2 × 10^-16^). **H)** Assembly member contribution (*P* = 3.53 × 10^-3^).

**Figure S3:**
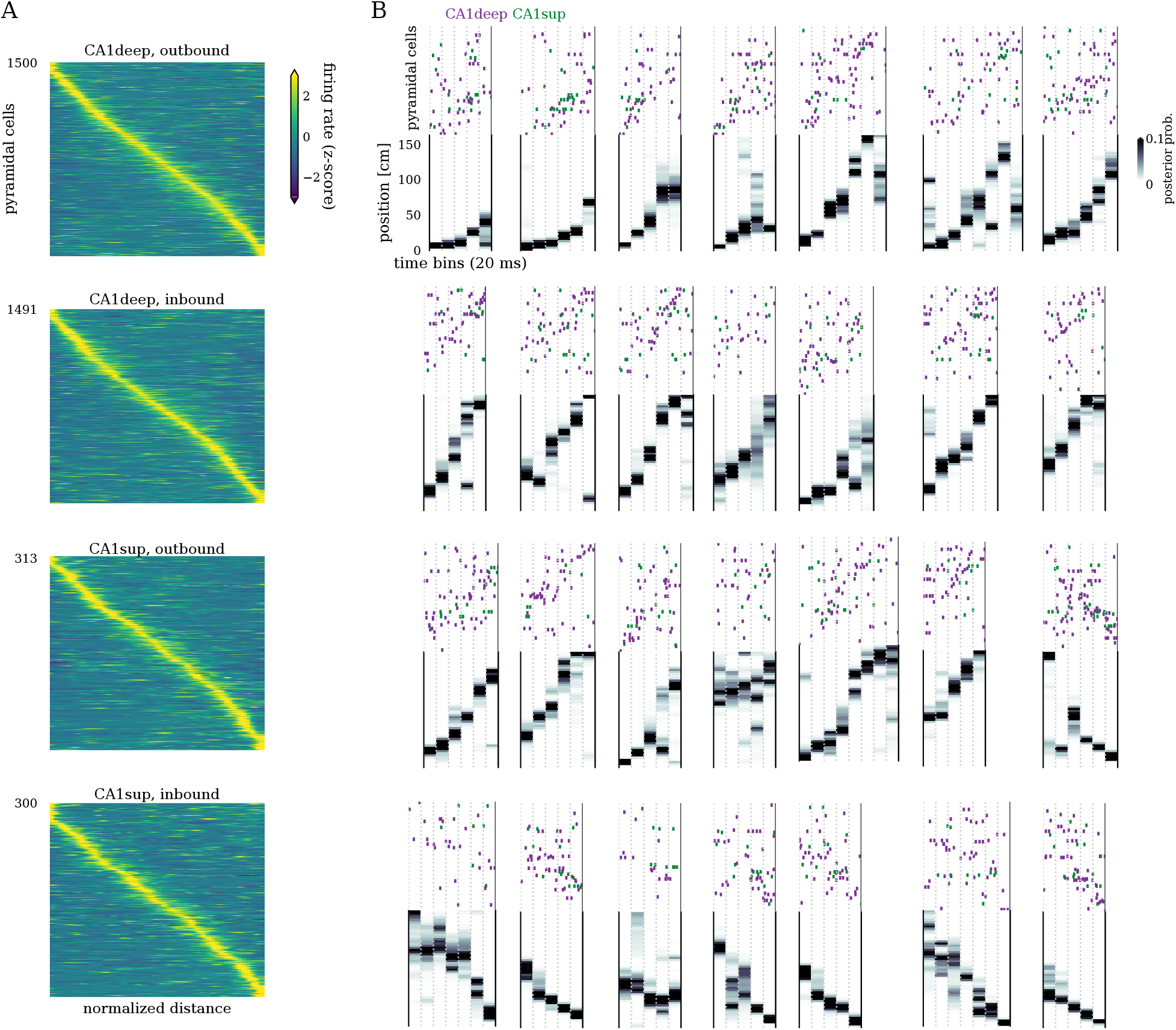
Example replay events. **A)** CA1deep and CA1sup place cells on the linear track in outbound and inbound directions. Each panel is sorted by the place field’s peak location. **B)** Example unit raster (top) during a SWR and decoded animal position for several replay events.

**Figure S4:**
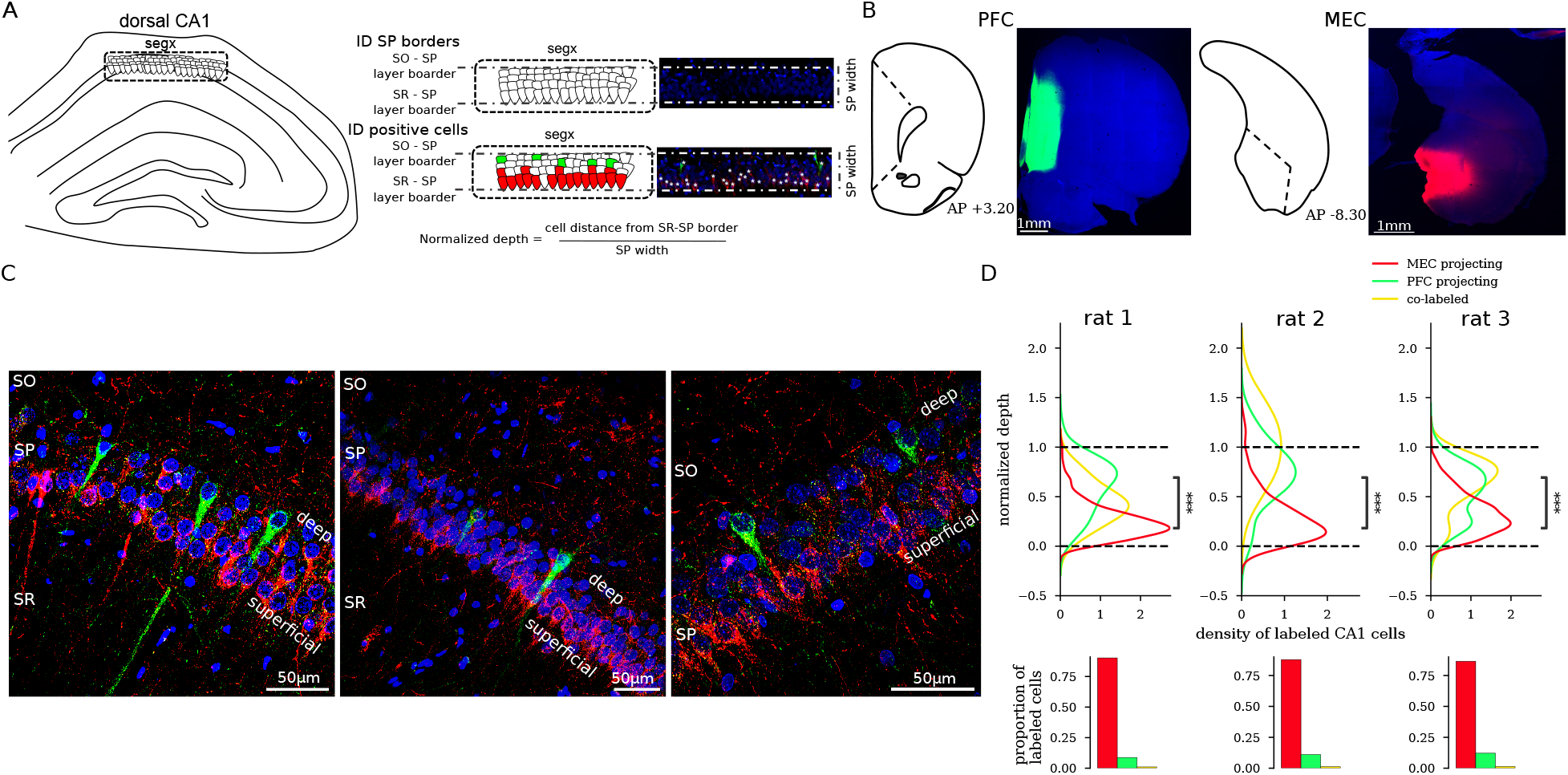
Retrograde tracing quantification of CA1 projecting to PFC and MEC. **A)** Left: Shows a diagram of how segments from the dorsal hippocampal CA1 pyramidal layer were defined. Right: shows the manual quantification process to estimate the cell’s relative distance from the pyramidal layer’s borders. **B)** Example histology of the injection sites in PFC and MEC in a different animal from Fig. 2. Blue: DAPI, Red: CTB-555, Green: CTB488. **C)** Example histology from different animals showing PFC projecting (green) and MEC projecting (red) cells. Blue: DAPI, Red: CTB-555, Green: CTB488. Scale bars: 50 *μm*. **D)** Quantification of soma location for labeled cells for individual rats (*P_s_* ≤ 5.84 × 10^-7^, rank-sum test, PFC-projecting versus MEC-projecting.) Bottom: proportion of labeled cells for each rat.

**Figure S5:**
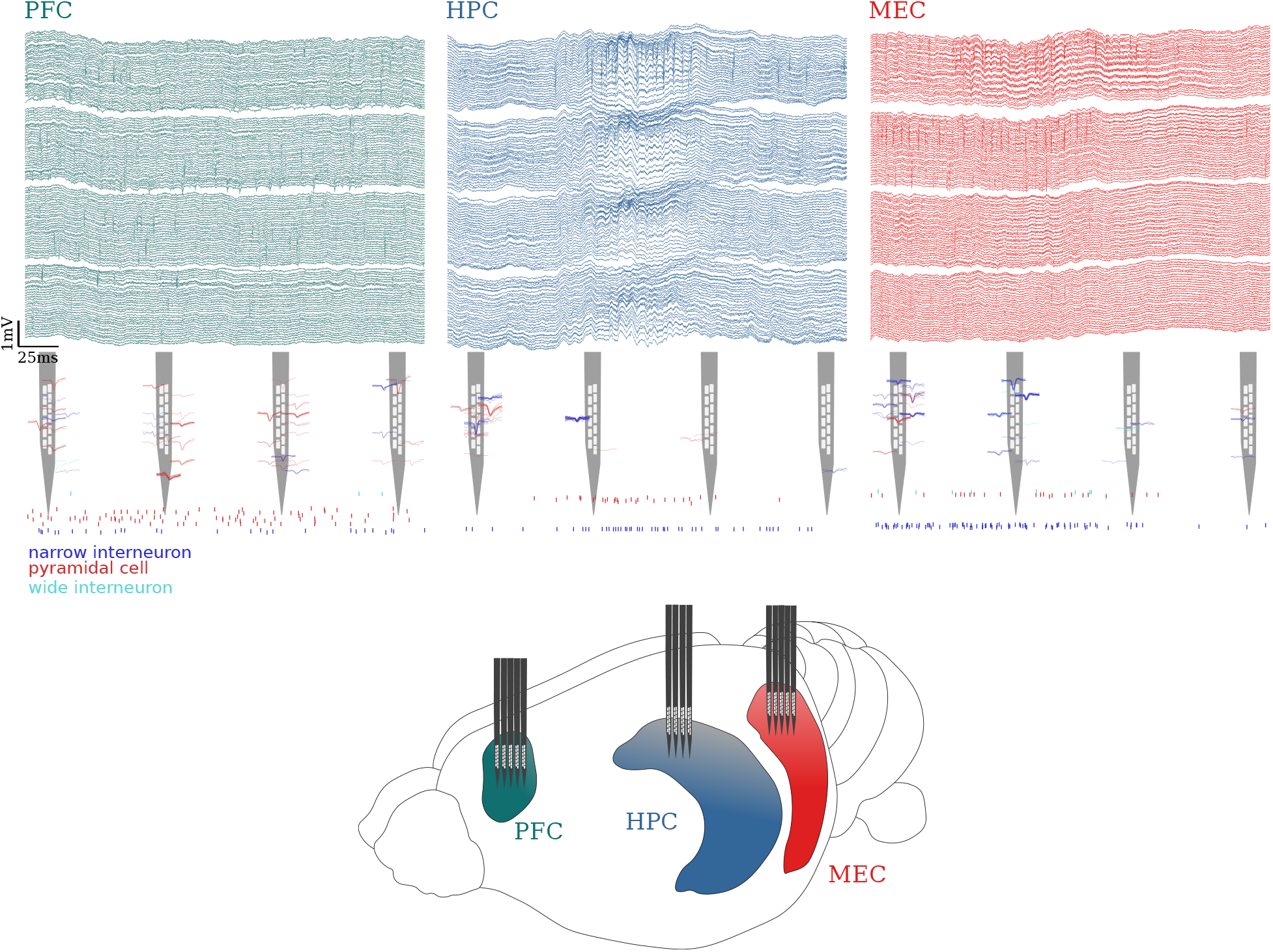
Depiction of multi-region recordings. Top: simultaneously recorded LFPs (wideband traces) from PFC, HPC, and MEC. Middle: spike waveforms from the current time window overlaid on a probe schematic and raster plot from pyramidal cells and interneurons. Bottom: schematic of multi-region recording in PFC, HPC, and MEC using three 128 channel 4 shank probes.

**Figure S6:**
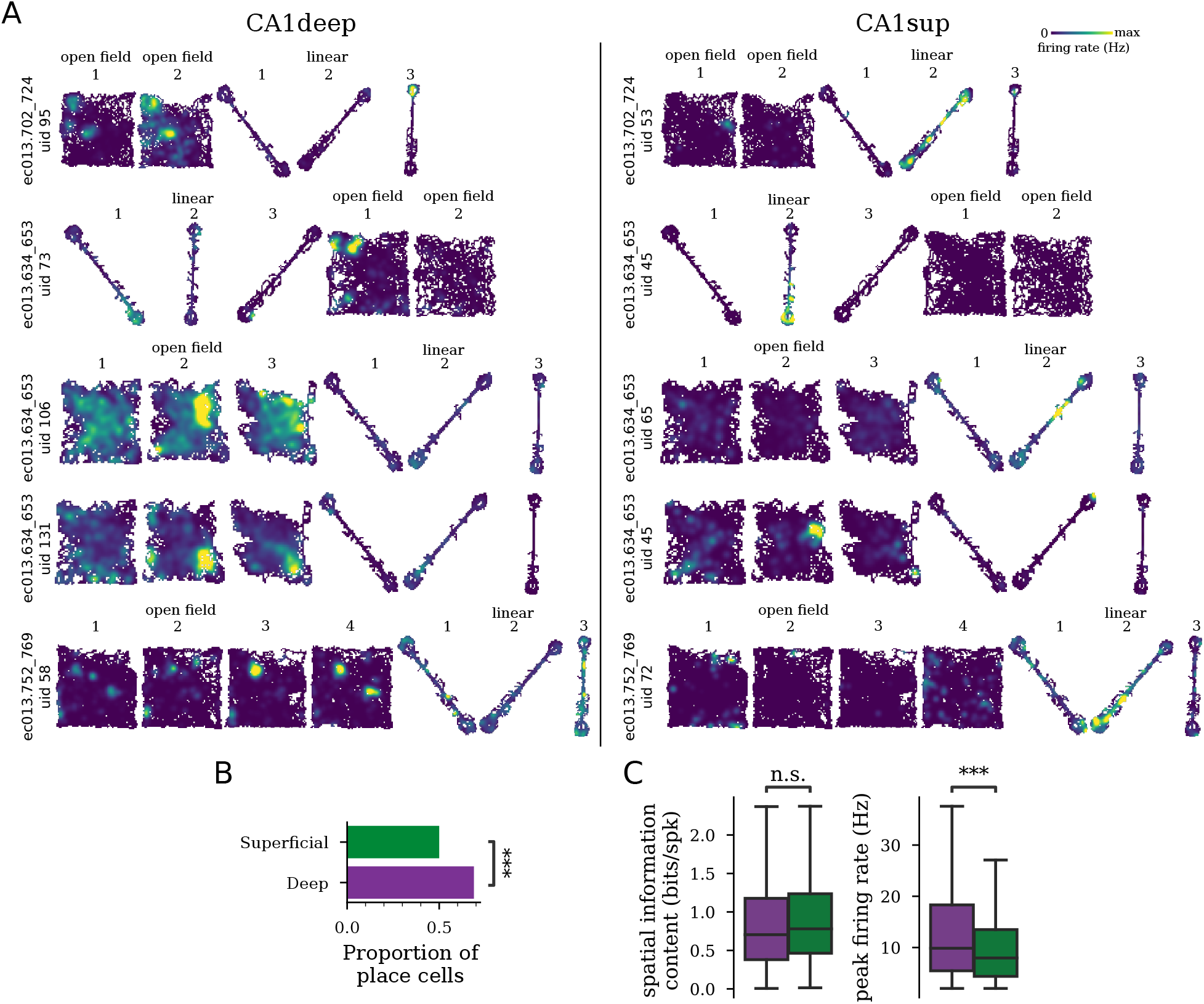
Place cells across multiple contexts. **A)** Example rate maps for five CA1deep and five CA1sup cells recorded across several contexts on the same day. Each row corresponds to a different cell. Darker colors indicate low firing rates and lighter colors indicate higher firing rates. Note how CA1deep express place fields and have elevated firing rates across multiple mazes, while CA1sup tends to be more selective. Color is scaled per individual cell. **B)** Proportion of cells with at least one place field in a recording session (CA1sup versus CA1deep *P* = 6.34 × 10^-15^, χ^2^). **C)** Spatial information content (*P* = 0.98, rank-sum test) and peak firing rate (*P* < 2 × 10^-16^).

**Figure S7:**
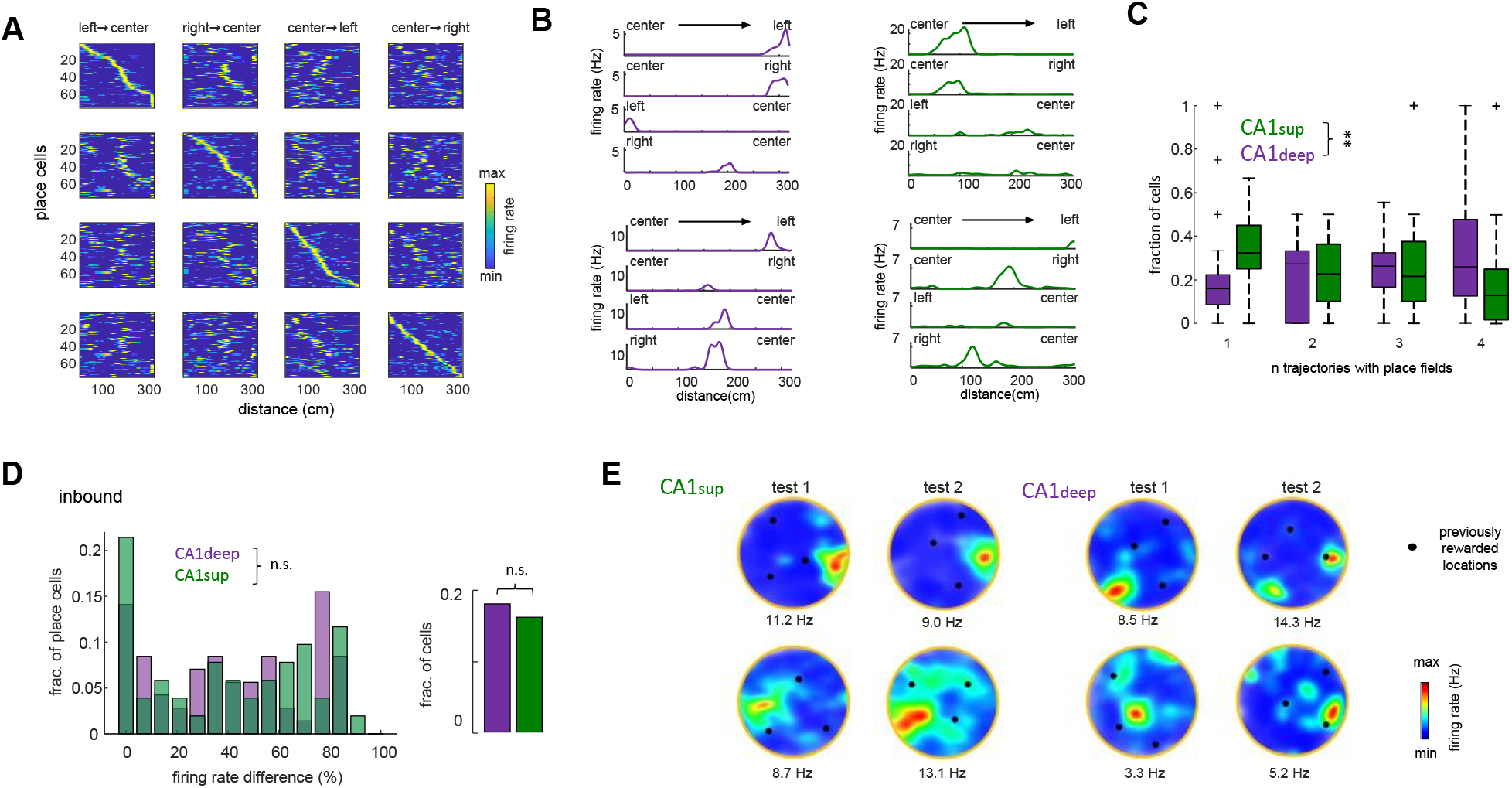
Place cells during memory tasks. **A)** Place cells in a representative session in the M-maze delayed alternation task. Each column corresponds to one of the four possible trajectories in the maze. In each row, cells were sorted according to the order of their place fields in one of the trajectories. This sorting resulted in a diagonal sequence of place fields for that trajectory and a random pattern for the others, evidencing the existence of unique place maps for each trajectory. In each subplot, rows correspond to individual place cell rate maps (normalized to the peak firing rate). **B)** Example of two CA1deep (left) and two CA1sup (right) place cell rate maps from the same session as A. **C)** Distribution of number of trajectories in which cells express place fields in the M-maze (n = 325 CA1sup and 296 CA1deep place cells, *P* = 0.0044, rank-sum test) **D)** Left: distribution of firing rate difference for left and right inbound trials for all cells with overlapping fields in the central arm (n = 71 CA1deep/ 51 CA1sup; *P* = 0.93, rank-sum test). Right: fraction of splitter cells (inset; *P* = 1, Fisher’s test). **E)** Examples of two CA1sup (left) and two CA1deep (right) place cells in the cheeseboard maze task. The left plot corresponds to test 1 session and the right plot to test 2 session. Black dots indicate reward location in the preceding learning session (during test rewards were not present). Note how CA1deep place cells remap in test 2 session towards new reward locations.

**Figure S8:**
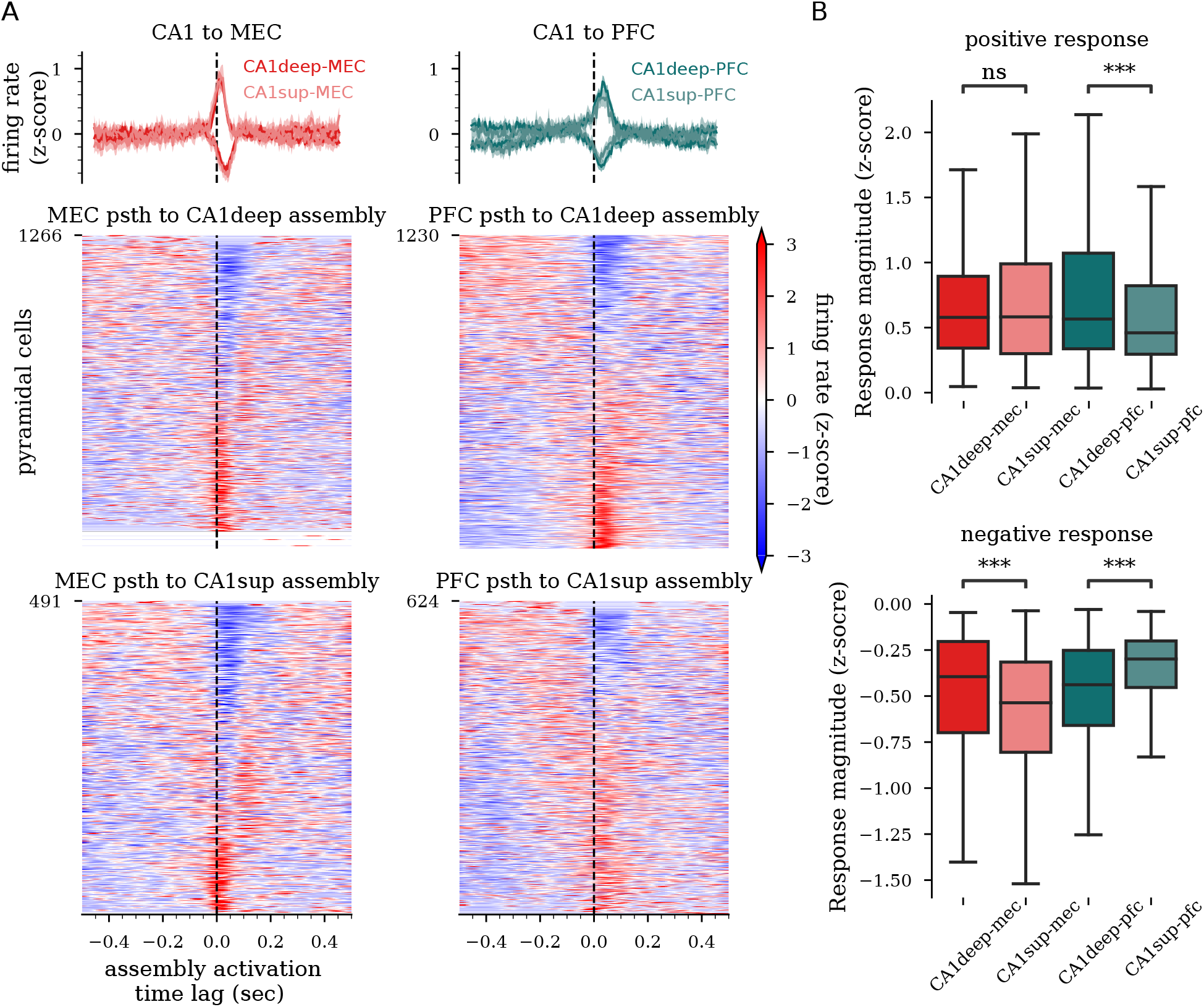
Cortical response of cortex to CA1 assembly activation in post-task sleep. **A)** Average (top) and individual unit (bottom) peri-assembly activation firing rate responses for CA1deep and CA1sup and MEC and PFC. **B)** Quantification of cortical single unit responses for CA1deep and CA1sup assembly activation. Units with positive (top) and negative responses are shown separately. *** *P* < 0.001 rank-sum test

